# Imidacloprid Movement into Fungal Conidia is Lethal to Mycophagous Beetles

**DOI:** 10.1101/2020.01.11.901751

**Authors:** Robin A. Choudhury, Andrew M. Sutherland, Mathew J. Hengel, Michael P. Parrella, W. Douglas Gubler

## Abstract

1. Applications of systemic pesticides can have unexpected direct and indirect effects on nontarget organisms, producing ecosystem-level impacts.
2. We investigated whether a systemic insecticide (imidacloprid) could be absorbed by a plant pathogenic fungus infecting treated plants and whether the absorbed levels were high enough to have detrimental effects on the survival of a mycophagous beetle. Beetle larvae fed on these fungi were used to assess the survival effects of powdery mildew and imidacloprid in a factorial design. Fungal conidia were collected from treated and untreated plants and were tested for the presence and concentration of imidacloprid.
3. Survival of beetles fed powdery mildew from imidacloprid-treated leaves was significantly lower than that of beetles from all other treatments.
4. Imidacloprid accumulated in fungal conidia and hyphae was detected at levels considered lethal to other insects, including coccinellid beetles.
5. Water-soluble systemic insecticides may disrupt mycophagous insects as well as other nontarget organisms, with significant implications for biodiversity and ecosystem function.

## Introduction

Systemic insecticides are considered valuable tools for pest management. Products containing water-soluble active ingredients are applied to roots via the substrate or to leaves and are readily translocated throughout the plant. Active toxins accumulate in the plant tissue and are toxic to arthropods feeding on the plants. Such applications protect active ingredients from ultraviolet light and other degradation factors, enabling them to persist longer in the environment. In addition, soil applications of systemic insecticides have sometimes been considered less harmful than sprays to beneficial insects active in the plant canopy while specifically targeting phytophagous pests (Pflueger & Schmuck, 1991).

Neonicotinoid insecticides, which disrupt insect nicotinic acetylcholine receptors, are commonly employed worldwide within pest management programs (Matsuda et al., 2005). Imidacloprid, the first commercially successful neonicotinoid, has excellent systemic properties. Imidacloprid products are labeled for use against many phytophagous pests in agricultural and urban landscapes, such as piercing/sucking insects, bark burrowers, and chewing beetle larvae, but activity is evident on a wide range of arthropods. Direct toxicity has been shown not only to common target pests such as aphids and whiteflies (Qu et al., 2015; Smith et al., 2016), but also to beneficial insects such as coccinellid beetles (Lee et al., 2017), hymenopteran parasitoids (Tappert et al., 2017), and predatory mites (Put et al., 2016).

A growing body of literature exists documenting the direct and indirect effects of imidacloprid and other neonicotinoids against pollinator insects (Easton & Goulson, 2013; Goulson, 2013). For instance, the imidacloprid-laced pollen from seed-treated sunflowers was implicated in deaths of foraging honeybees and decreases in honey production in France (Bonmatin et al., 2003). Guttation fluids from treated plants have also been identified as routes to exposure for nontarget arthropods (Hoffmann & Castle, 2012). Indirect toxicity to beneficial insects may also readily occur. It has been shown that coccinellid predators may ingest lethal doses of imidacloprid through the consumption of sessile homopteran prey that have ingested this systemic material through phytophagy (Grafton-Cardwell & Gu, 2003). Even consumption of honeydew from homopterans feeding on treated plants can negatively impact the fecundity and survival of beneficial insects (Calvo-Agudo et al., 2019). Imidacloprid and its plant metabolites move readily through various organisms at different trophic levels, imparting toxicity to susceptible insects. Such trophic movement violates the integration clause of integrated pest management.

Plant pathogenic fungi that utilize plant water and nutrients, such as the powdery mildews (Erysiphales), may act as reservoirs for systemic insecticides, and therefore may be toxic to susceptible arthropods if consumed. The cosmopolitan coccinellid tribe Halyziini is known to obligately consume Erysiphales conidia and hyphae, providing ecosystem services in natural and agricultural settings throughout the world (Sutherland & Parrella, 2009).

Using this model, we sought to determine whether imidacloprid can move into fungal hyphae and whether this movement would negatively affect beetles that fed on the contaminated hyphae.

## Materials and methods

Adults of the halyziine coccinellid beetle *Psyllobora vigintimaculata* (Coleoptera: Halyziini) were collected from a vineyard in Fresno, CA in fall 2011. Beetles were reared in a growth chamber (PGR-15, Conviron Ltd., Winnipeg, Canada) that was kept at 25°C and 50% relative humidity under fluorescent lights with a 14-h photoperiod on pumpkin plants (*Cucurbita pepo* cv. Sorcerer) infected with the cucurbit powdery mildew pathogen, *Podosphaera xanthii* (syn. *Podosphaera fusca*). To encourage a uniformly aged population for our bioassay, we introduced approximately 400 beetles of mixed sex into a separate growth chamber containing pumpkin plants infected ten days prior. After four days of egg deposition, adults were removed from the colony and egg broods left to hatch.

Concurrently, pumpkin seeds were planted with Sunshine Mix potting medium 1 (SunGro Horticulture, Bellevue, Wash.) into five 6-inch pots in trays and flood irrigated with either 17.4% imidacloprid (EPA Registration# 264-763-AA-67760) or deionized water in a greenhouse. After ten days, one tray each of imidacloprid-treated and untreated plants were removed from the greenhouse, inoculated with *P. xanthii* (by gently transferring conidia from infected leaves of other plants), and placed into a growth chamber (25°C, 50% RH). The other two trays were kept uninoculated in the greenhouse (20° ± 10°C, 50% ± 20% RH).

Fifteen days later, leaves from all four seedling groups were removed and cut petioles inserted into 2 ml glass vials with deionized water, sealed with parafilm. These vials were then placed into inverted transparent plastic quart (~ 950 ml) containers and sealed. One first- or second-instar beetle larva was introduced to each container. Containers with uninfected leaves (without powdery mildew) were supplemented with one 15 mm leaf disk cut from untreated infected plants, to provide a food source for *P. vigintimaculata* larvae, known as obligate mycophages (Davidson, 1921). Excised leaves and vials were changed every four days to prevent wilting; supplemental infected leaf disks were changed every three days. In this way, treatments included: 1) untreated and uninfected with infected leaf disc supplement: IM-/PM-, 2) untreated and infected with powdery mildew: IM-/PM+, 3) imidacloprid-treated and uninfected, with infected leaf disc supplement: IM+/PM-, and 4) imidacloprid-treated and infected with powdery mildew: IM+/PM+. Each treatment was represented by 15 replicate containers, for a total of 60 bioassay arenas (see Figure S1). Containers were maintained at ambient room temperature (22°C) and 12h photoperiod. Mortality, moribundity, and developmental stage were assessed for beetle larvae daily until emergence of adults from pupae. Moribund larvae that were immobile for two or more days were judged dead and removed from the assay. Observations continued until all larvae were dead or had successfully pupated. Conidia from both imidacloprid treated and untreated leaves were vacuum aspirated into a microcentrifuge tube and tested for imidacloprid concentration using liquid chromatography and tandem mass spectrometry (see supplemental materials for specific methods used).

Survival analyses were conducted in R v. 3.5.2 using the ‘survival’ package (Therneau, 2015). To analyze the main effects of imidacloprid and powdery mildew treatments on survival, in addition to the interaction effects, we generated both a full regression model and performed a pairwise logrank test using Bonferoni-adjusted *p*-values.

## Results

Qualitative examination of survival curves suggests that the treatment groups began to diverge by ~24h post exposure (Fig. 1), and that 100% of beetles raised on powdery mildew from imidacloprid-treated leaves died before the end the study. Survival regression; including imidacloprid treatment, powdery mildew treatment, and the interaction between the two as explanatory factors; revealed that the effects of imidacloprid treatment was substantially higher [hazard ratio (HR)= 3.66, 95% confidence interval (CI): 1.86–7.18] than the effect of powdery mildew food availability (HR=2.18, 95% CI: 1.13–4.19). We also found a strong interaction between the imidacloprid treatment and the presence of powdery mildew (HR=4.99, 95% CI: 1.23–20.21).

**Fig. 1:**
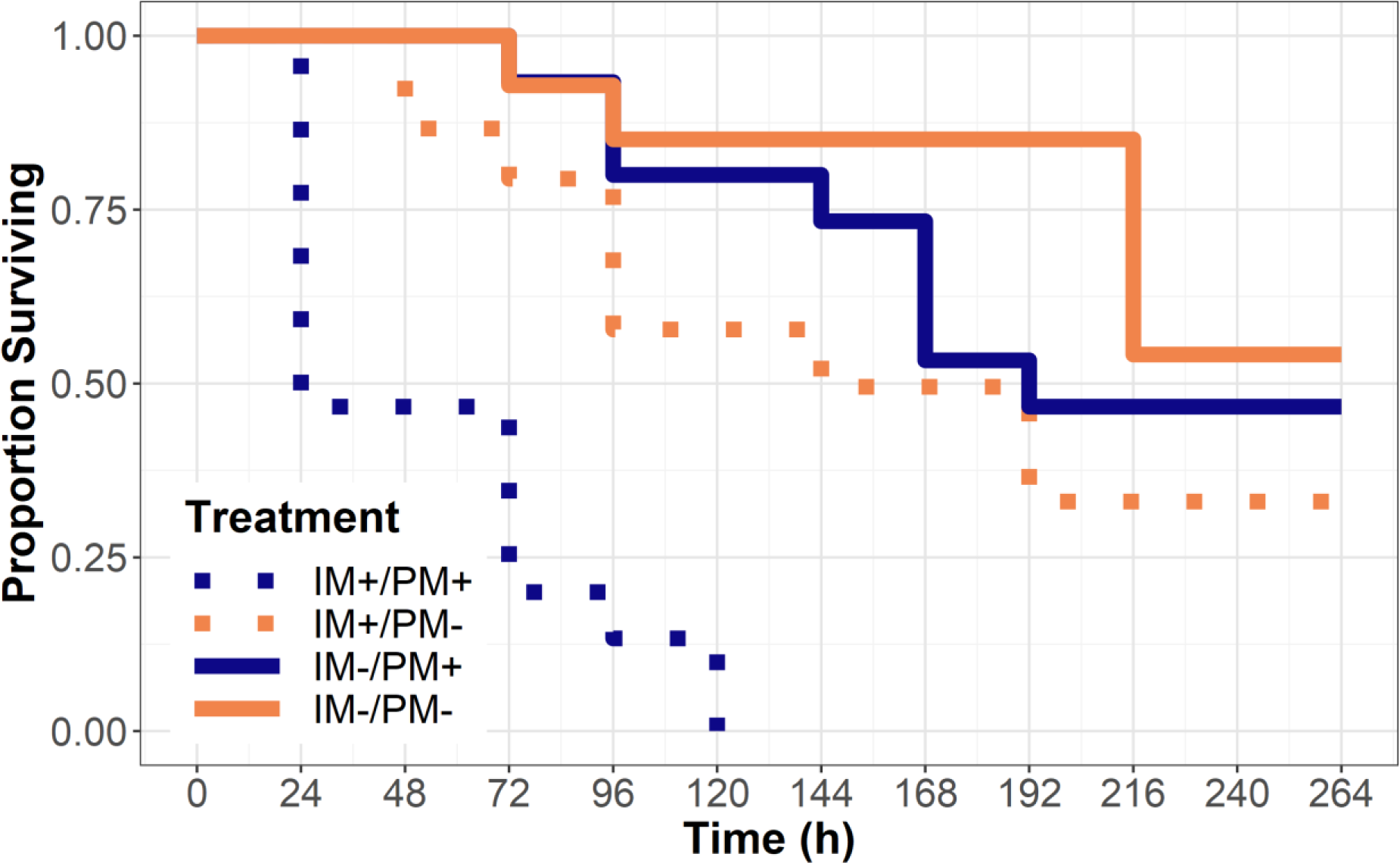
Survival of obligately mycophagous beetle larvae, *P. vigintimaculata*, after being confined to leaves from plants either treated or untreated with imidacloprid (IM+/IM-) and either infected or uninfected by powdery mildew fungus *Podosphaera xanthii* (PM+/PM-). Larvae confined to uninfected leaves (PM-) were regularly provided sustenance by way of supplemental leaf discs from infected and untreated plants.

Chemical analysis detected increased levels of imidacloprid in conidial samples taken from leaves excised from plants treated with imidacloprid. Powdery mildew conidia collected from leaves treated 15-days prior had imidacloprid concentrations of 33.6 µg/g (n=3), whereas conidia from untreated leaves exhibited concentrations below the detectable level (0.05 µg/g) (n=1) (Figure S2). Conidia collected from leaves 22 days after the imidacloprid treatment exhibited a mean concentration of 7.65 µg/g (n=5), while untreated leaves continued to have concentrations below the detectable level (n=4).

## Discussion

This study represents the first record of the tri-trophic movement of imidacloprid from plant to fungus to insect, ultimately creating significant indirect mortality of a nontarget organism and potentially disrupting a pathway for biological control of an important group of pathogens. We observed that all beetle larvae fed on powdery mildew grown on imidacloprid-treated plants perished within 120 hours. In contrast, larvae confined to imidacloprid-treated leaf arenas but regularly provided with leaf discs from untreated infected plants exhibited no significant differences in survival as compared to those fed on powdery mildew grown on untreated plants. These observations, as well as the survival regression results, suggest that the insecticide did not impact survival of larvae through volatilization from treated leaves nor through incidental phytophagy. Instead, mortality was associated strictly with consumption of fungal structures growing from treated plant tissue. This conclusion is supported by previous observations that soil-applied imidacloprid is readily translocated into various tissues of *C. pepo* plants (Stoner & Eitzer, 2012) and that lethal effects of imidacloprid consumption on three different coccinellid species has been observed at 6.03 µg/g (Krischik et al., 2015). In general, imidacloprid is considered toxic to coccinellid larvae, with lethal residues at concentrations as low as 2.6 µg/g (Mizell & Sconyers, 1992).

Mycophagous beetles may play an important role in management and detection of powdery mildew disease, and reductions in their populations might exacerbate powdery mildew disease outbreaks in susceptible crops. Sutherland and Parrella (2006) found that a single larvae of *P. vigintimaculata* removed an average of 6.3 cm2 of leaf area affected by powdery mildew during development from egg to adult. Furthermore, Peduto et al. (2011) found that incidence of *P. vigintimaculata* adults was positively correlated with incidence of early season grapevine powdery mildew disease, suggesting a possible use of *P. vigintimaculata* as a bioindicator. The use of mycophagous beetles in greenhouse settings to directly consume powdery mildew or as early indicators of disease may help direct management of diseases in such controlled settings (Parrella & Lewis, 2017). Mycophagous beetles from the Halyziini are observed throughout the world in association with powdery mildew infections (Sutherland & Parrella, 2009), suggesting that the potential for indirect mortality of mycophagous beetles after exposure to systemic insecticides is widespread.

The ecosystem-level impacts of direct and indirect exposure to lethal doses of systemic insecticides is still being explored. Several studies have explored the tri-trophic movement of systemic insecticides and the subsequent disruption of natural predators and parasitoids (Calvo-Agudo et al., 2019; Grafton-Cardwell & Gu, 2003; Lee et al., 2017; Put et al., 2016; Tappert et al., 2017). While some studies demonstrated the lethal impacts of direct applications of pesticides to mycophagous beetles (Lee et al., 2017; Sutherland et al., 2010), our study showed that indirect exposure to imidacloprid through the fungal food source can also rapidly lead to death in mycophagous beetles. Widespread measurements of reductions in insect diversity and abundance have recently been attributed to intensive agricultural activities and associated pesticide inputs (Sánchez-Bayo & Wyckhuys, 2019); it’s possible that trophic movement of water-soluble toxins, as observed in our study, play a part in this global problem.

In the future, exploring natural populations of *Psyllobora* species located near fields that had been treated with imidacloprid might help to clarify the ecological role of imidacloprid in this tri-trophic interaction.

## Supporting Information

Additional Supporting Information may be found in the online version of this article under the DOI reference: xxxxx

### File S1

Online supporting material.

