## SUPPLEMENTAL FILES for "Imidacloprid Movement into Fungal Conidia is Lethal to Mycophagous Beetles"

**Online Supporting Information**

**Supplementary Materials**

*Liquid Chromatography / Tandem Mass Spectrometry of Samples:*

Materials. Imidacloprid (CAS Registry No. 138261-41-3, 98%) was obtained from Bayer Crop Science, Kansas City, MO. All solvents and reagents were pesticide grade or better. Water was prepared using a Milli-Q reagent water system. Specifications for filtration are cited below.

Preparation of Standard Solutions. A Stock solution (1.00 mg/mL) of imidacloprid was prepared by adding 10 mg (corrected for purity) of the analytical grade compound to a 10 mL volumetric flask and brought up to volume with acetonitrile (MeCN). Fortification solutions were prepared by diluting aliquots of the stock solution and diluting up with MeCN, resulting in 100, 10, 1, 0.1 and 0.01 μg/mL solutions. Calibration solutions for LC-MS/MS analysis were prepared by taking various volumes of the fortification solutions and diluting up to volume in water/acetonitrile (50:50, v/v) resulting in calibration standards over a range of 0.1–200 pg/μL. Standard solutions were stored at ~ 5 °C and were stable for 1 month.

Extraction. Typically, a 5 mg aliquot of powdery mildew hyphae was weighed into a 1.5 mL disposable micro centrifuge tube (recovery samples were fortified at this point) and 0.5 mL of MeCN was added. The sample was vortexed briefly and then shaken vigorously for 1 minute. The sample was vortexed again and then centrifuged for 2 min. at 6400 RPM. A 250 µL aliquot of the supernatant was taken and diluted with 250 µL of water. If necessary, the diluted sample was filtered through a 0.2 µm Teflon syringe filter (National Scientific, Rockwood, TN). Typically, unknown samples were diluted to a final sample volume of 1 mL (equivalent to 5 mg of powdery mildew hyphae per mL), such that the LOQ for imidacloprid would be equivalent to 0.25 pg/µL.

Sample Analysis**.** Sample analysis was conducted with an Agilent 1200 series autosampler and binary pumps coupled to an Agilent 6430 tandem mass spectrometer via an electrospray ionization source (ESI) (Agilent Technologies, Wilmington, DE). The ESI source was operated in positive ionization mode with 10 L/min of drying nitrogen gas at 300 °C. The nebulizer was set to 50 psi of nitrogen and the capillary was operated at 4500 volts. The mass spectrometer was operated in multiple reactant monitoring mode (MRM). See **Table S1** for compound conditions. Chromatographic separation was accomplished with an Agilent Zorbax SB-$C_{18}$ column (30 × 32.1 mm ID, 3.5 μm particle size). The mobile phase composition was 68:32 (v/v) 0.05% formic acid in water/acetonitrile with a flow rate of 200 μL/min (isocratic). Injection volume was 10 μL. Sample residues were quantified using a linear standard curve method with 1/x weighting. Typically, the primary transition was used for quantitation and the secondary transition and ion ratio were used for compound confirmation.

**Table S1:** Compound specific information for chromatography and mass spectrometry conditions.

| Compound | Primary Transition | Secondary Transition | Ion Ratio 2°/1° (%)[a] | $r^2$ | RT (min) |
|---|---|---|---|---|---|
| Imidacloprid | 256 → 209 | 256 → 175 | 99 | 0.99 | 0.96 |

[a] Ion ratio determined by abundance of secondary transition divided by the abundance of primary transition · 100%.

**Table S2:** Average imidacloprid recoveries from powdery mildew hyphae.

| | Fortification Level (µg/g) | | |
|---|---|---|---|
| Compound | 0.05[a] (n=4) | 5.0[a] (n=5) | 50[a] (n=3) |
| Imidacloprid | 91 ± 12 | 90 ± 5 | 89 ± 1 |

[a] Values are mean percent recovered ± standard deviation; n is the number of replicates.

**Table S3:** non-parametric comparisons for each treatment pair using Wilcoxan methods. Asterisks represent significance at the Bonferroni adjusted p-value of 0.0083.

| | PM+IM+ | PM+IM- | PM-IM+ | PM-IM- |
|---|---|---|---|---|
| PM+IM+ | ---- | <0.0001* | 0.0014* | 0.0002* |
| PM+IM- | ---- | ---- | 0.133 | 0.7655 |
| PM-IM+ | ---- | ---- | ---- | 0.1348 |
| PM-IM- | ---- | ---- | ---- | ---- |

**Figure S1**: experimental setup of a portion of the imidacloprid-*Psyllobora* bioassay, including detached cucurbit leaves inside of sealed polyethylene containers and *Psyllobora* beetles on the surface of the leaves.

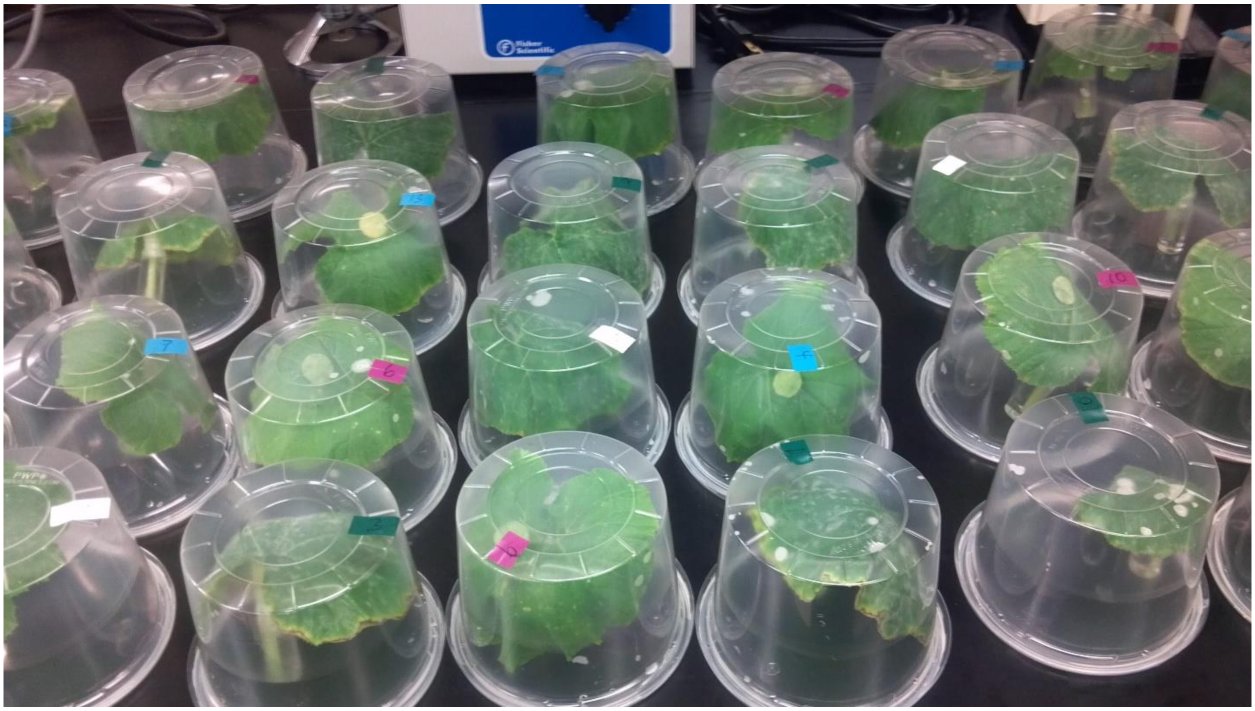

**Figure S2**: Concentration (ppm) of imidacloprid in powdery mildew conidia collected from treated (IM+) and untreated (IM-) plants 15 and 22 days after treatment. Points represent individual samples, and the overlaying boxplots represent the quantiles.

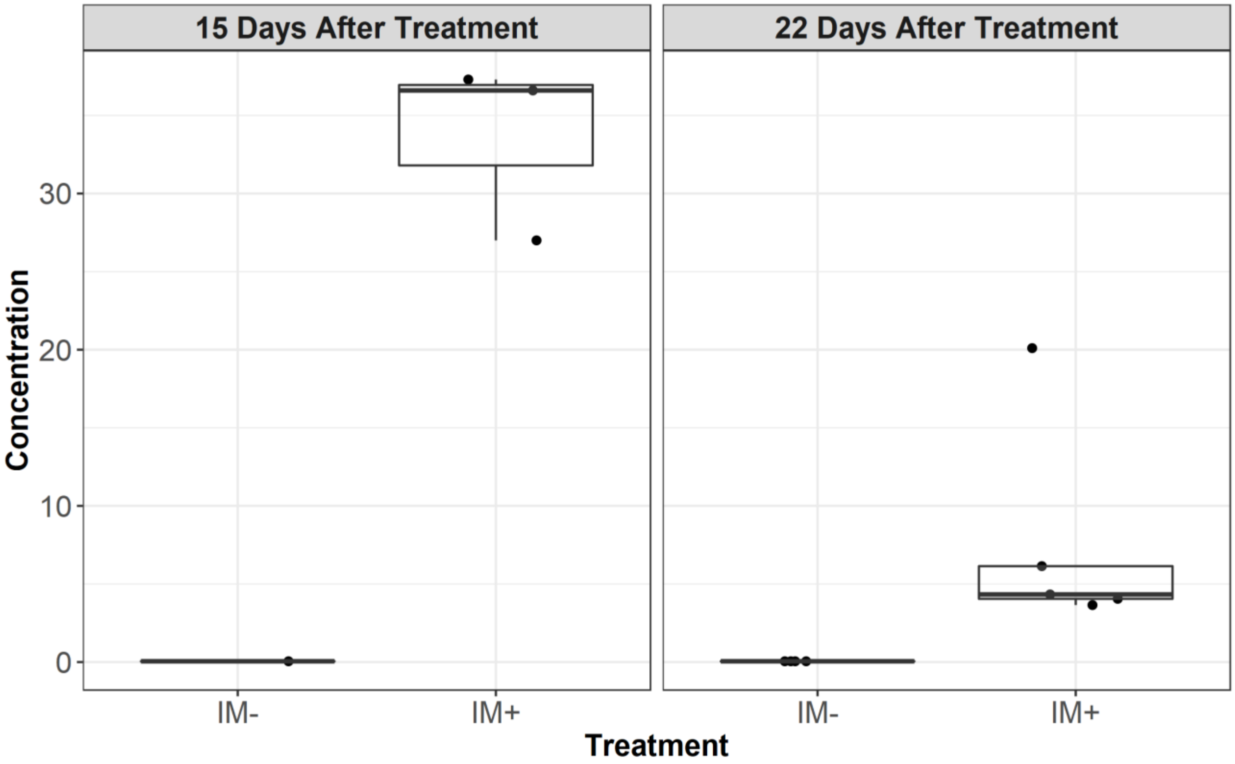
